# Novel Internalin P homologs in *Listeria*

**DOI:** 10.1101/2022.01.19.476994

**Authors:** Kayla N Conner, Joseph T Burke, Janani Ravi, Jonathan W Hardy

**Author notes:** Co-primary authors, contributed equally (Kayla N Conner, Joseph T Burke).

## Abstract

*Listeria monocytogenes* (*Lm*) is a bacterial pathogen that causes listeriosis in immunocompromised individuals, particularly pregnant women. Several virulence factors support the intracellular lifecycle of *Lm* and facilitate cell-to-cell spread, allowing it to occupy multiple niches within the host and cross protective barriers, including the placenta. One family of virulence factors, internalins, contributes to *Lm* pathogenicity by inducing specific uptake and conferring tissue tropism. Over 25 internalins have been identified thus far, but only a few have been extensively studied. Internalins contain leucine-rich repeat (LRR) domains which enable protein-protein interactions, allowing *Lm* to bind host proteins. Notably, other *Listeria* species express internalins but cannot colonize human hosts, prompting questions regarding the evolution of internalins within the genus *Listeria*. Internalin P (InlP) promotes placental colonization through interaction with the host protein afadin. Though prior studies of InlP have begun to elucidate its role in *Lm* pathogenesis, there remains a lack of information regarding homologs in other *Listeria* species. Here, we have used a computational evolutionary approach to identify InlP homologs in additional *Listeria* species. We found that *L. ivanovii londoniensis* (*Liv*) and *L. seeligeri* (*Ls*) encode InlP homologs. We also found InlP-like homologs in *L. innocua* and the recently identified species *L. costaricensis*. All newly identified homologs lack the full-length LRR6 and LRR7 domains found in *Lm*’s InlP. These findings inform on the evolution of one key *Lm* virulence factor, InlP, and serve as a springboard for future evolutionary studies of *Lm* pathogenesis as well as mechanistic studies of *Listeria* internalins.

**Impact Statement:** The intracellular bacterial pathogen *Listeria monocytogenes* can breach protective barriers in the pregnant host, allowing for the colonization of the placenta in pregnant women and resulting in numerous adverse pregnancy outcomes. Previous studies aimed at delineating the mechanisms behind placental colonization of *L. monocytogenes* identified a key virulence factor, internalin P (InlP). The internalin family of proteins has been studied extensively due to their conservation in the *Listeria* genus and their contribution to virulence and pathogenicity in *L. monocytogenes*. Still, many questions remain regarding the evolution of internalins and their potential roles in non-pathogenic *Listeria*. Our work addresses this gap in knowledge by 1) identifying additional InlP homologs in *Listeria*, including *L. ivanovii, L. seeligeri, L. innocua, and L. costaricensis*, and 2) characterizing these homologs using computational evolutionary methods to compare their primary sequences, domain architectures, and structural models. Together, our findings contribute to the field by providing insights into the evolution of one key member of the internalin family as well as serving as a catalyst for future studies of InlP and its role in *Listeria* pathogenesis.

## Introduction

Prenatal infection remains a major public health concern. Annually, nearly 13 million infants are born prematurely worldwide, and an estimated 30% of these preterm births can be attributed to prenatal infection, though the actual number may be higher due to the subclinical nature of many prenatal infections^1^. To better detect and treat these infections to prevent adverse pregnancy outcomes, we must better understand the pathogens that cause them. *Listeria monocytogenes* is widely used in prenatal infection research due to its well-characterized lifecycle and ease of use in laboratory experiments^2,3^.

The *Listeria* genus comprises 17 species, including the human pathogens *L. ivanovii* (*Liv*) and *L. monocytogenes* (*Lm*)^4^. These Gram-positive facultative intracellular bacterial pathogens are the most typical causative agent of listeriosis in humans^4,5^. While relatively rare, listeriosis can result in severe morbidity and mortality in immunocompromised individuals^5,6^. Pregnant women are particularly at risk for listeriosis, as *Lm* can colonize the placenta and cause adverse pregnancy outcomes such as preterm birth, neonatal meningitis, miscarriage, and stillbirth^5^. *Lm* employs several virulence factors that aid in its invasion of various host niches and the breach of protective host barriers^5,6^. Previous studies have addressed the roles of various *Lm* virulence factors, such as ActA, Internalin A (InlA), and Internalin B (InlB) in the context of pregnancy^7,8^. Faralla *et al*. followed up with two studies focusing on Internalin P (InlP), a key virulence factor for the invasion of the placenta^9,10^.

Typically, listeriosis begins with the consumption of contaminated food items. Once in the digestive system, *Lm* uses several virulence factors, including the internalins, to colonize gut epithelial cells and spread throughout the host^5,6,11^. Internalins contribute to this spread by conferring tissue tropism; for example, InlA binds E-cadherin on gut epithelial cells while InlB binds C-Met expressed by hepatocytes^11^. These interactions are enabled by Leucine-Rich Repeat (LRR) domains found in all internalins^11,12^. LRR domains are found in an array of functionally diverse proteins across the domains of life, such as ribonuclease inhibitors in eukaryotes (humans, pigs) and virulence factors in prokaryotes (*Yersinia pestis, L. monocytogenes)*^12^. Internalins may have as few as four (InlG) or as many as fourteen (InlA) LRR domains^5,11^. Notably, internalins and other virulence factors are relatively well-conserved across the genus, including species that are considered non-pathogenic to humans, such as *L. seeligeri* and *L. innocua*^4,13,14^. While the details remain unclear, differences in pathogenicity have been attributed to minor genetic variations and differences in the expression of virulence factor genes^15^. The precise roles of the various internalin genes in *Listeria and their* evolutionary relationships remain critical open questions in *Listeria* biology.

Internalin P (InlP) is an *Lm* virulence factor known to enhance placental colonization in the pregnant host. This is likely accomplished by enabling *Lm* to transcytose through the basal membrane underlying the syncytiotrophoblast, the protective outer layer of placental cells that serves as a barrier between maternal and fetal blood^9^. Further characterization revealed that InlP encompasses nine LRR domains and binds the human protein afadin, which is a nectin-like protein found in cell-cell junctions and thought to play a significant role in cellular adhesion^10^.

Initial InlP studies identified a structural homolog of InlP, Lmo2027, in *Lm*, but information regarding InlP homologs in other *Listeria* species has been incomplete^9^. In this study, we used comparative genomics and protein sequence-structure-function analyses to identify InlP homologs in the genomes of *L. seeligeri* (*Ls*), *L. ivanovii londoniensis* (*Liv*), *L. innocua* (*Lin*), and *L. costaricensis* (*Lc*) (**Fig. 3, Table S1**). The bioinformatic analysis presented here serves as a springboard for future studies of *Listeria* evolution and pathogenesis pertaining to the internalin protein family, including its ability to colonize the human placenta.

## Materials and Methods

### Identification of InlP Homologs

To identify InlP_*Lm*_ homologs across evolutionary lineages, we submitted the InlP_*Lm*_ amino acid sequence (accession: WP_014601135.1) to MolEvolvR (http://www.jravilab.org/molevolvr)^16^. The query returned hits for homologous proteins across bacterial phyla. While many species carried homologous proteins (*e*.*g*., *Nostoc spp*. and *Beggiatoa leptomitoformis*), we chose to filter out hits with low similarity and divergent domain architectures and genomic contexts for our detailed study; we thus focused on the *Listeria* genus (including 45,530 *L. monocytogenes*, 740 *L. innocua*, 169 *L. seeligeri*, 44 *L. ivanovii*, and 1 *L. costaricensis* genomes). Within this dataset, we selected the hits with the highest percent similarity and unique domain architectures as representative homologs for further analysis (**Figure 1; Table S1**). Accession numbers provided by MolEvolvR were used to query the NCBI RefSeq Protein Database for corresponding nucleotide sequences and locus tags for homologous genes^17^. BioCyc^18^ (https://biocyc.org) and NCBI RefSeq^17^ protein databases were used to identify genomic contexts (neighboring genes).

**Figure 1:**
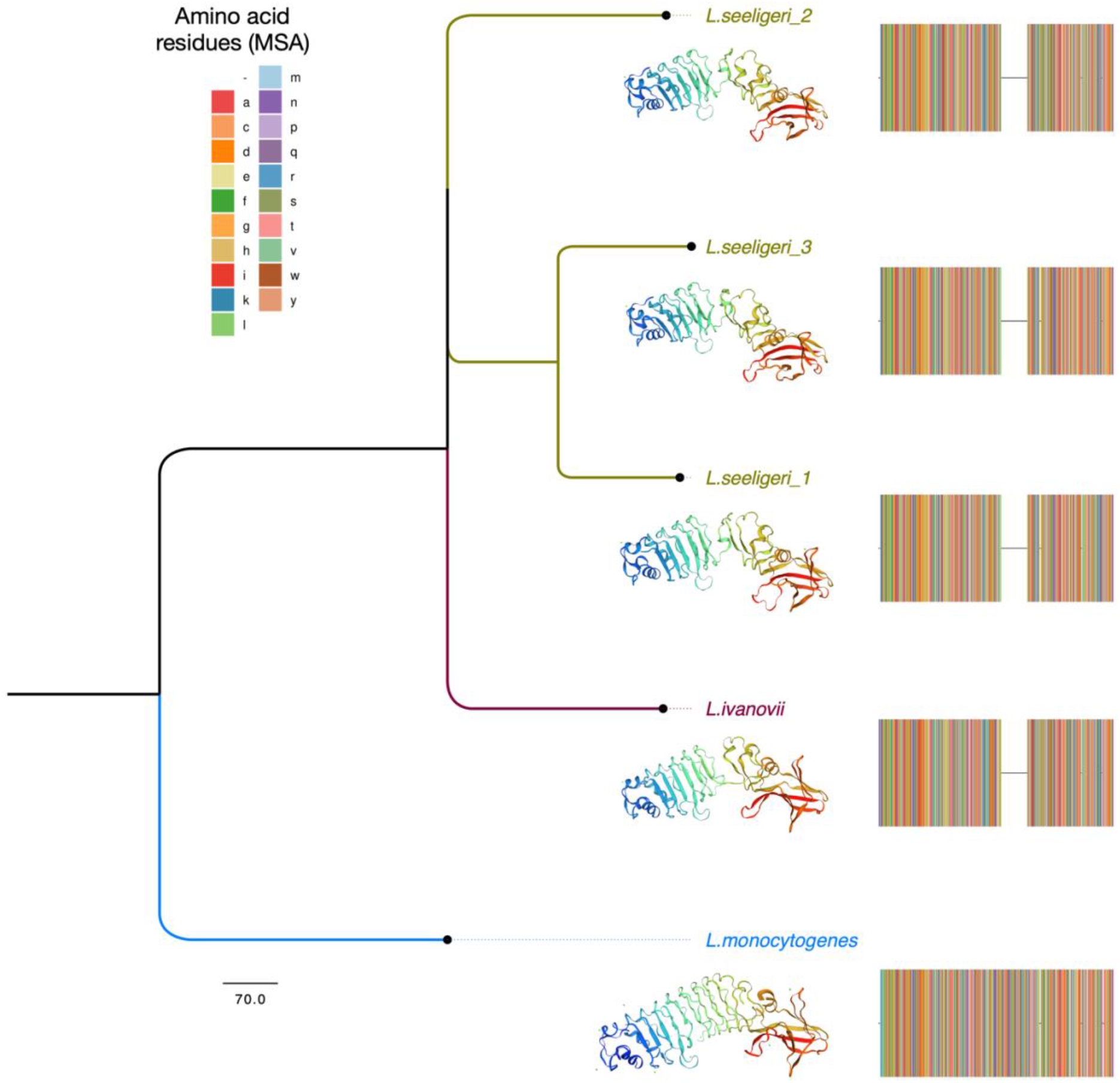
Phylogeny and Structure Models of InlP and Representative Homologs. A phylogenetic tree was generated using the amino acid sequences of InlP homologs in *L. monocytogenes* (blue), *L. ivanovii londoniensis* (Purple), and *L. seeligeri* (green). Three-dimensional models were generated using SWISS-MODEL with the crystal structure for InlP_*Lm*_ (PDB: 5hl3) as a template. Multiple sequence alignments were generated using MolEvolvR and illustrate the complete LRR6 and LRR7 insertion present in InlP_*Lm*_. The legend shows the colors of the amino acid residues indicated in the multiple sequence alignment

### Calculation of Percent Identity and Percent Similarity

Percent identity and percent similarity values for predicted homologs were provided by MolEvolvR^16^ (**Table S1**; https://github.com/jravilab/inlp_listeria). Nucleotide sequences for *inlP* in *L. monocytogenes, L. ivanovii londoniensis, L. seeligeri*, and *L. costaricensis* were aligned using Clustal Omega^19^ (https://www.ebi.ac.uk/Tools/msa/clustalo/), and resulting alignments were submitted in FASTA format to the Sequence Manipulation Suite^20^ (SMS; http://www.bioinformatics.org/sms2) to calculate percent nucleotide identity. The homolog similarity and identity matrix was generated using MatGAT2.01 with the BLOSUM 62 matrix and default options^21^.

### Multiple Sequence Alignment, Phylogenetic Trees, and Protein Models

Multiple sequence alignments for homologous amino acid sequences were generated using Kalign^24^ and visualized using JalView (Version 2.11.1.4)^25^. Phylogenetic trees were constructed using Kalign^24^, FastTree, and FigTree. Three-dimensional protein models were produced using SWISS-MODEL with the *L. monocytogenes* Internalin P crystal structure (PDB: 5hl3) as a template^26,27^. All data, analyses, and visualizations for InlP Listeria homologs are available here: https://github.com/jravilab/inlp_listeria.

## Results

### *Listeria ivanovii londoniensis* and *Listeria seeligeri* encode Internalin P Homologs

To begin investigating evolutionary conservation of the *L. monocytogenes* Internalin P (InlP_*Lm*_), we started with an extensive homology search and protein characterization of InlP-like proteins in diverse lineages across the tree of life using MolEvolvR^16^ (http://jravilab.org/molevolvr). Most homologs were present only within the genus *Listeria*. To further ensure that all homologs are being identified, we picked other representative InlP homologs from *L. ivanovii* and *L. seeligeri* as new starting points for our homology search and characterization (using MolEvolvR^16^; see *Methods*). We found several hits in our multi-start search including proteins that resembled other Internalins (*e*.*g*., InlB), and others that carried neither the signature Internalin_N or LRR domains characteristic of internalins (**Fig. 2 and S1**). Therefore, we restricted our full set of homologs to only InlP-like proteins resulting in 64 representative proteins with distinct domain architectures from each *Listeria* species including *L. monocytogenes, L. seeligeri, L. ivanovii, L. innocua*, and *L. costaricensis* (**Fig. 2 and S1; Table S1**). Homologs from *L. seeligeri* (*Ls*) and *L. ivanovii* (*Liv*) showed >65% amino acid similarity compared to InlP_*Lm*_, while homologs from *L. innocua* (*Lin*) and *L. costaricensis* (*Lc*) showed <35% and 55% similarity, respectively (**Fig. 3**). We found that several homologs lacked predicted signal peptide domains suggesting they are not secreted like InlP; this was further corroborated by the presence of predicted transmembrane LPXTG motifs, which indicate that these homologs are more likely to be membrane-anchored InlB-like proteins rather than secreted InlP homologs (**Fig. 2 and S1**). Investigation of the InlP-like proteins in *Lc and Lin* showed that it is unlikely that they are functional InlP homologs (discussed below). Further investigation of domain architectures and genomic contexts of putative homologs ultimately revealed one InlP homolog encoded within the *L. ivanovii londoniensis* genome and three InlP homologs encoded by *L. seeligeri* (discussed below). Here, we refer to these homologs as InlP_*Lm*_, InlP_*Li*_, InlP_*Ls1*_, InlP_*Ls2*_, and InlP_*Ls3*_, respectively, to indicate species and gene order.

**Figure 2:**
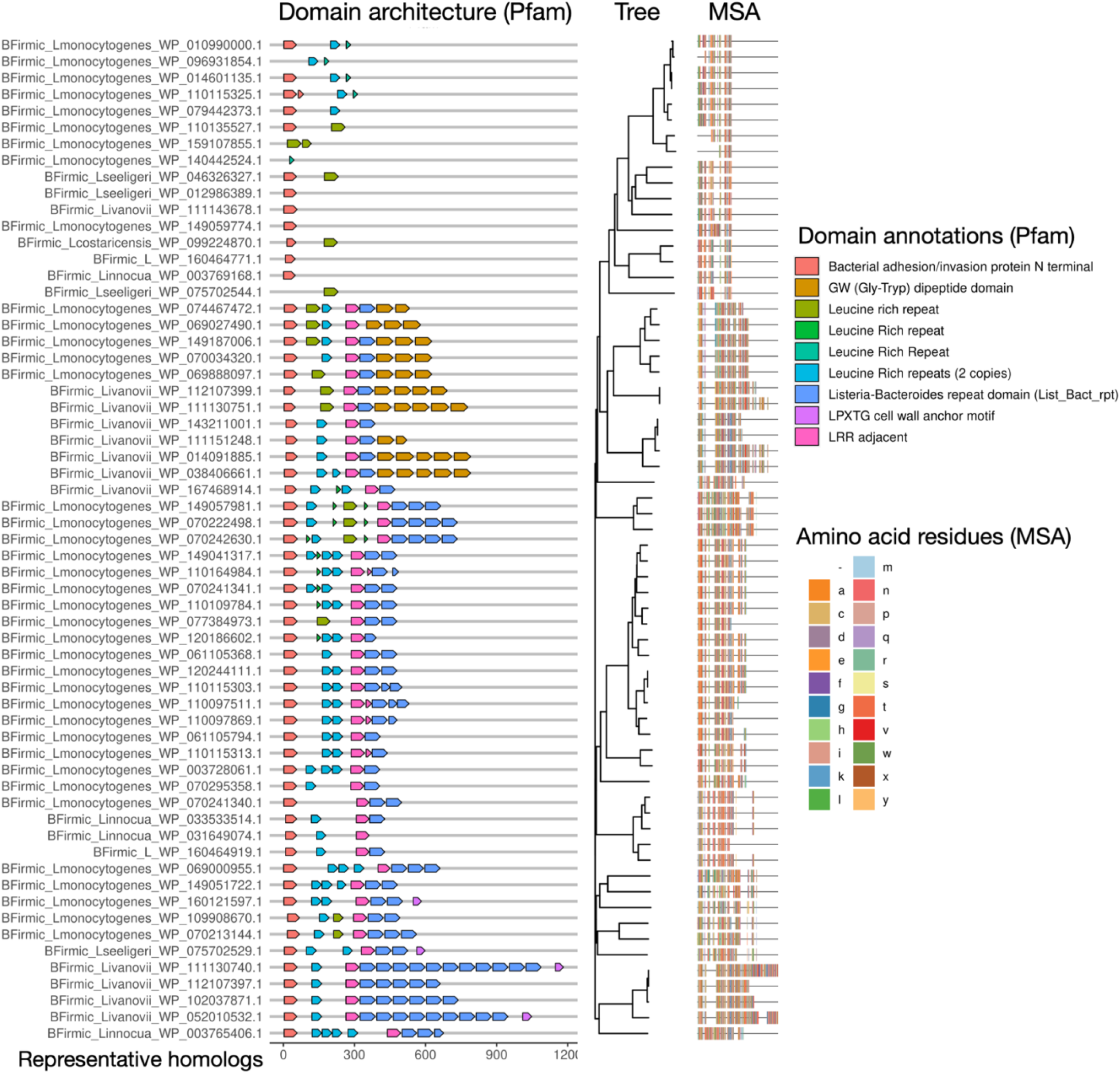
Phylogeny and Domain Architectures of Putative Internalin P Homologs. A multiple sequence alignment and phylogenetic tree were generated using the amino acid sequences of putative InlP homologs identified using MolEvolvR and five *Listeria* InlP starting points (See *Methods*). The phylogeny of InlP-like proteins has been overlaid with their domain architectures (generated using Pfam database, MolEvolvR). The legend shows the colors of the amino acids indicated in the multiple sequence alignment and Pfam domain annotations.

**Figure 3:**
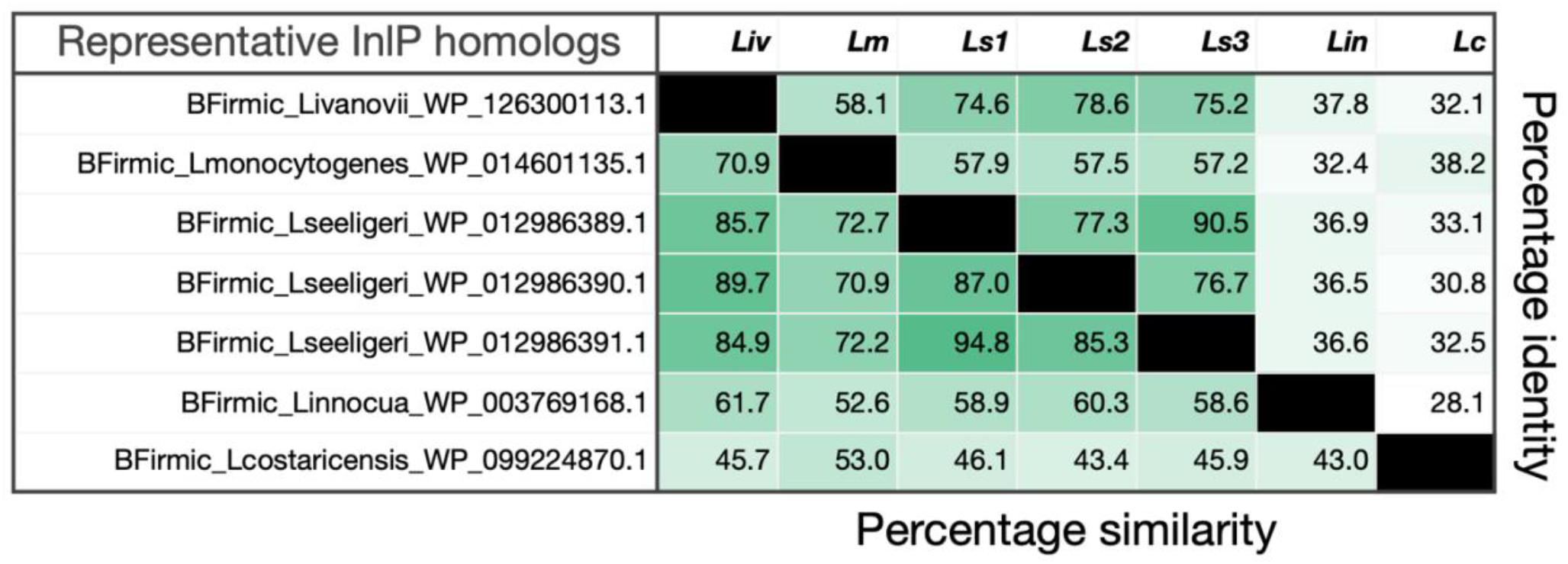
Percent Similarity and Identity of Internalin P Homologs in representative *Listeria* species. Percent similarity and percent identity were calculated for the amino acid sequences for representative Internalin P homologs from five *Listeria* species, *L. monocytogenes, L. ivanovii, L. seeligeri, L. costaricensis*, and *L. innocua*. Matrix showing similarity and identity values was generated using MatGAT2.01 with the BLOSUM 62 matrix and default options selected.

We determined the similarity of InlP homologs in *Liv* and *Ls* at the nucleotide and amino acid levels towards functional characterization. The *inlP* gene in *L. ivanovii londoniensis* (*inlP*_*Liv*_), as well as the three *L. seeligeri* paralogs (*inlP*_*Ls1*_, *inlP*_*Ls2*_, and *inlP*_*Ls3*_), shared ∼70% identity with the *inlP*_*Lm*_ gene and ∼52–65% identity at the amino acid level when compared to InlP_*Lm*_ (**Fig. S2**). However, the newly identified homologs in *L. ivanovii londoniensis* and *L. seeligeri* shared much higher percent amino acid similarity with InlP_*Lm*_ — InlP_*Liv*,_ InlP_*Ls1*,_ and InlP_*Ls3*_ showed ∼70% similarity to InlP_*Lm*_ (**Fig. 3**). InlP_*Ls2*_ showed a striking 94% similarity to the *L. monocytogenes* InlP (**Fig. S2**). Notably, the flanking *L. seeligeri* paralogs, InlP_*Ls1*_ and InlP_*Ls3*_, were more similar to each other than to the third paralog or to InlP_*Lm*_ (**Fig. 3**). To further investigate these new *Listeria* InlP proteins, we next explored their genomic neighborhoods.

Because the *Listeria* genus maintains a high degree of synteny across species, we investigated the genomic contexts of identified homologs compared to the InlP_*Lm*_ gene, which is flanked upstream by an amino acid permease gene and downstream by an NADPH dehydrogenase gene (**Fig. 4A**). We hypothesized that functional homologs of InlP_*Lm*_ would be flanked by these same genes in other *Listeria* species. We, therefore, determined the genomic neighborhoods of *inlP* homologs in *L. ivanovii londoniensis, L. ivanovii ivanovii, L. seeligeri, L. costaricensis*, and *L. innocua* (*see Methods*). We found that the *inlP* genes in *L. ivanovii londoniensis* and *L. seeligeri* were flanked upstream by an amino acid permease gene and downstream by an NADPH dehydrogenase gene mirroring the *L. monocytogenes* genomic context, suggesting the identified homologs are likely true homologs of the *inlP* gene (**Fig. 4**). Interestingly, we found that the gene encoding the InlP-like protein in *Lc* was flanked upstream by an amino acid permease and downstream by a gene encoding a LapB repeat-containing protein, inconsistent with genomic neighborhoods seen in other *Listeria* species (**Fig. 4E**). Also inconsistent with other genomic neighborhoods, the InlP-like protein in *Lin* was flanked upstream by a DUF5110-containing protein-encoding gene and downstream by the *ssrA* gene (**Fig. 4F**). To quantify the similarity of the flanking genes in *L. ivanovii* londoniensis, *L. seeligeri*, and *L. monocytogenes*, we calculated their pairwise similarity. At the amino acid level, the products of these flanking genes had >95% similarity to *L. monocytogenes* (**Fig. S3**). Notably, while *L. ivanovii londoniensis* encoded an *inlP* homolog in this region, *L. ivanovii ivanovii* did not (**Fig. 4**). Additionally, while *L. ivanovii londoniensis* encoded only one copy of *inlP* (EL212_RS12905; *inlP*_*Li*_), *L. seeligeri* encoded three copies (LSE_RS12040, LSE_RS12045, and LSE_RS12050; *inlP*_*Ls1*,_ *inlP*_*Ls2*,_ and *inlP*_*Ls3*,_ respectively) (**Fig. 4**).

**Figure 4:**
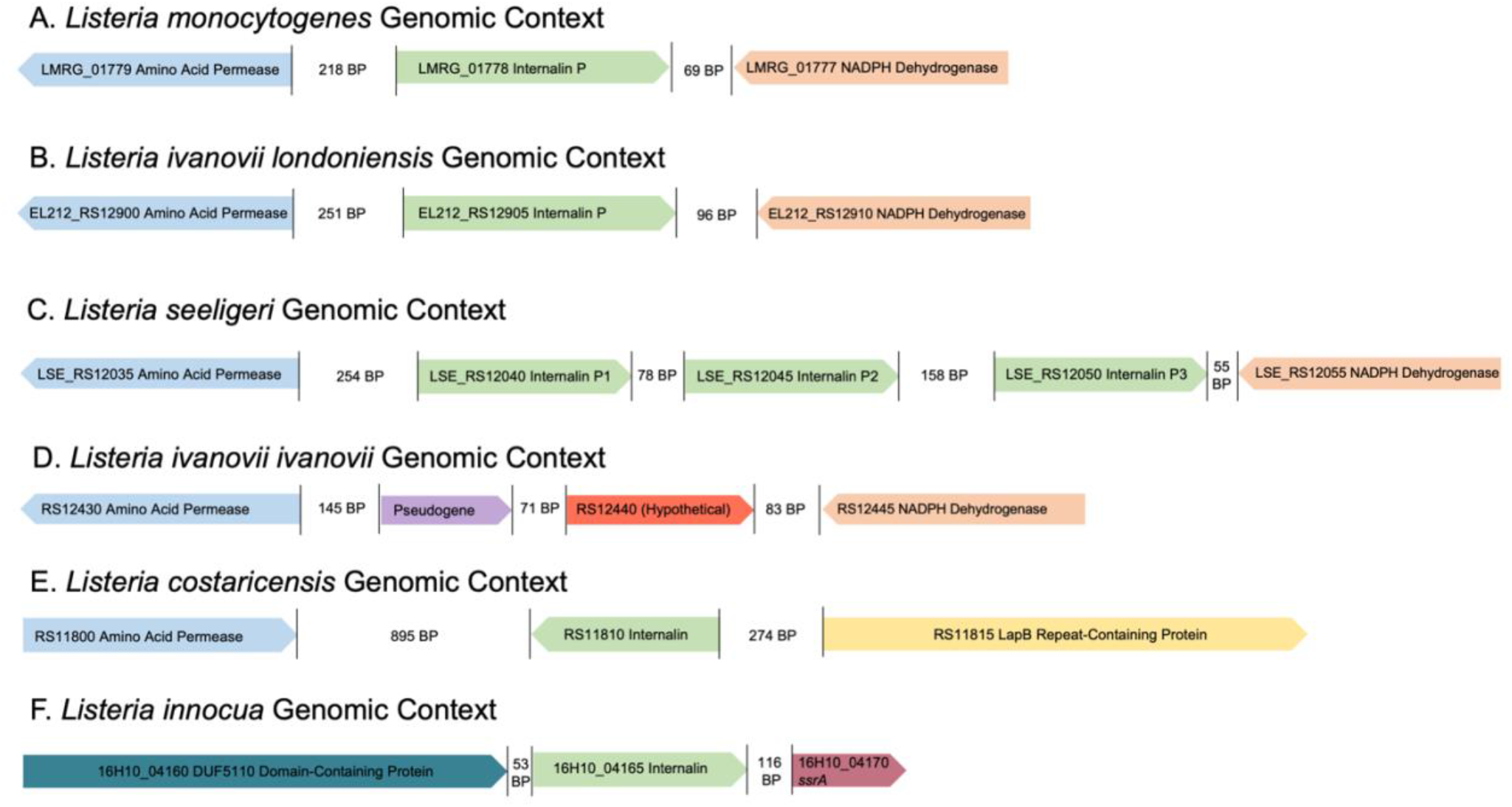
Genomic Context of Newly Identified Internalin P Gene Homologs. Genes homologous to the *L. monocytogenes inlP* (A) were identified in various other *Listeria* species. Gene order was maintained in *L. monocytogenes, L. seeligeri*, and *L. ivanovii londoniensis*. Notably, *L. seeligeri* encodes three copies of the *inlP* gene. All homologs categorized as “true” homologs were flanked upstream by an amino acid permease gene (blue) and downstream by an NADPH dehydrogenase gene (orange). *L. ivanovii ivanovii* contains a pseudogene (purple) and an uncharacterized gene encoding a hypothetical protein (red) in this region. Putative homologous genes in *L. costaricensis* and *L. innocua* did not mirror genomic neighborhoods seen in the other *Listeria* species. All *inlP* homologs are represented in green. Genomic context was determined using RefSeq genomic records and the BioCyc genome browsers for each species (see *Methods*).

### Internalin P Homologs in *L. ivanovii* and *L. seeligeri* lack the full-length LRR6 and LRR7 domains found in *L. monocytogenes* InlP

To delineate the evolution of InlP within *Listeria*, we generated a multiple sequence alignment (**Fig. 5**) and constructed a phylogenetic tree of the homologs (**Fig. 1**). While we observed several amino acid substitutions throughout the length of the proteins, the most striking difference between InlP_*Lm*_ and its homologs was in the Leucine-Rich Repeat (LRR) regions — a partial lack of LRR6 and complete lack of LRR7 — in the *L. ivanovii* and *L. seeligeri* homologs (**Fig. 1**). Additionally, we noted a lack of conservation in a previously described calcium-binding loop^10^ present in InlP_*Lm*_ (amino acid residues 132– 135; **Fig. 1**). This observation was of particular interest since this calcium-binding loop might play a role in protein signaling or stabilization of protein-protein interactions between InlP_*Lm*_ and host afadin. To better visualize the structural differences in these homologs, we generated models of InlP_*Li*_, InlP_*Ls1*_, InlP_*Ls2*_, and InlP_*Ls3*_ based on the previously resolved crystal structure of InlP_*Lm*_ (**Fig. 1**; see *Methods*). These models illustrate the similarity in the overall structure of the five homologous proteins, and the lack of LRR7 and full-length LRR6 are visible in *Li* and *Ls* homologs (**Fig. 1**; green/yellow regions). Additionally, the calcium-binding loop region is discernible in all five homologous proteins but appears structurally diverse in *L. ivanovii* and *L. seeligeri* homologs.

**Figure 5:**
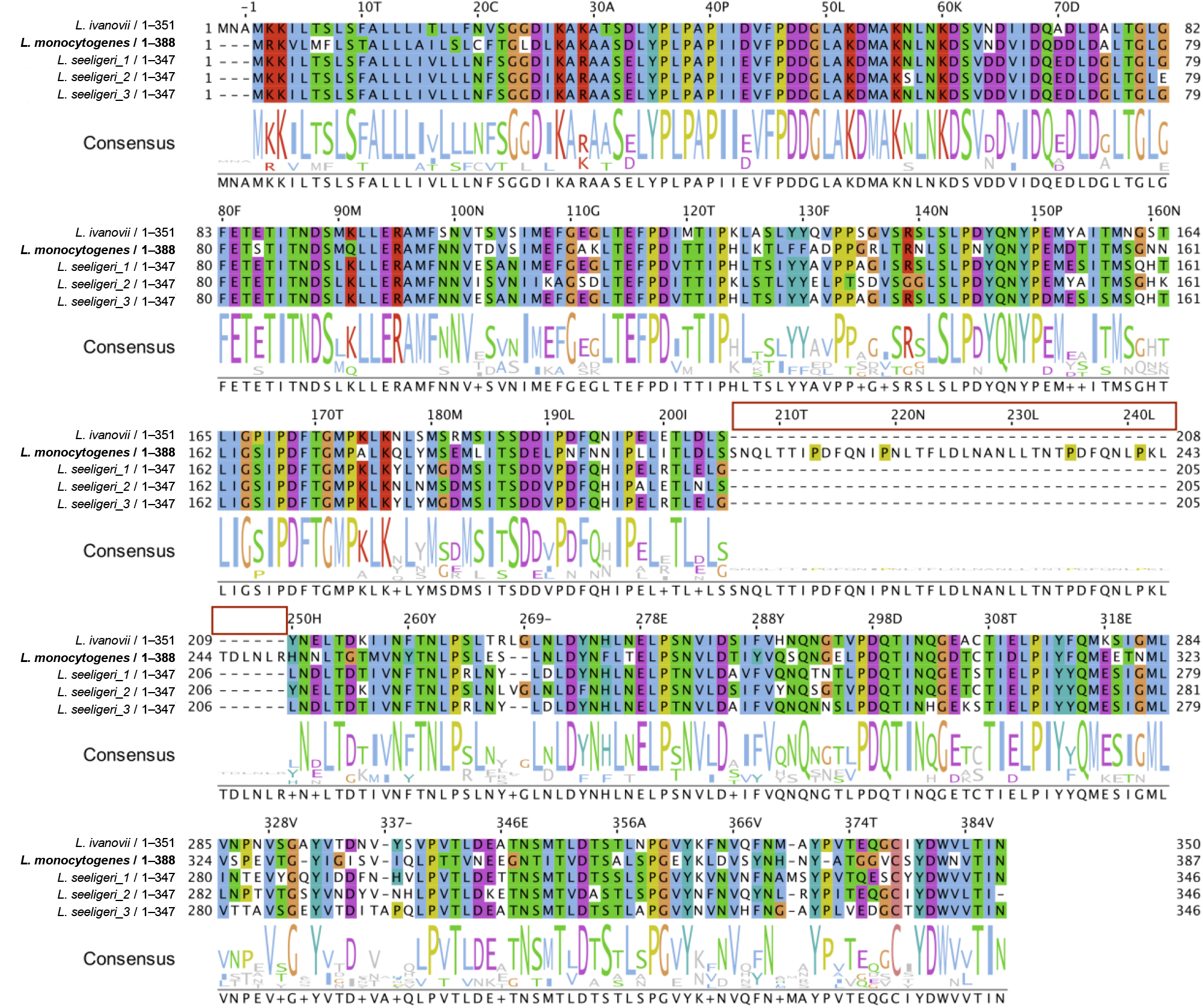
Multiple Sequence Alignment of Internalin P Homologs. Amino acid sequences for identified Internalin P homologs in *L. monocytogenes, L. ivanovii londoniensis*, and *L. seeligeri* were aligned using Kalign and visualized using Jalview. Below the alignment is the consensus sequence for the four homologs. The red box indicates the insertion present only in the InlP_*Lm*_ homolog. The *L. monocytogenes* homolog is used as the reference InlP protein (for residue numbering).

In summary, we have discovered novel Internalin P homologs in *Listeria*, traced their evolution, and uncovered potential functional implications pertaining to heterogeneity in key InlP domains. All InlP homolog data (along with characterizations in terms of domain architectures and modeling) are available at https://github.com/jravilab/inlp_listeria.

## Discussion

While previous studies have addressed many of the physical and mechanistic properties of InlP, the conservation of the *inlP* gene within or outside of the *Listeria* genus remains incompletely characterized. Here, we have provided insight into Internalin P in other *Listeria* species aside from *L. monocytogenes* that will drive future mechanistic studies of InlP as well as evolutionary studies of *Listeria* pathogenesis pertaining to the internalins.

First, we analyzed the InlP amino acid sequence with MolEvolvR^16^ to retrieve homologous proteins across evolutionary lineages. MolEvolvR^16^ is a powerful new bioinformatic web application to characterize protein families using molecular evolution and phylogeny (http://jravilab.org/molevolvr). The MolEvolvR^16^ InlP search returned a list of potential homologs including those found in *Listeria* species. In this article, we focus on homologs in *Listeria* since these species carried the classic Internalin and LRR domains. Using MolEvolvR^16^, we identified InlP-like proteins in *L. innocua, L. seeligeri, L. ivanovii*, and *L. costaricensis*; only homologs in *L. seeligeri* and *L. ivanovii* londoniensis expressed an amino acid percent similarity value >65%. We found that the lower identity proteins are more likely to be Internalin B homologs based on their sequence, domain architecture, and structure.

To determine if the newly identified proteins were true homologs of InlP_*Lm*_, we explored their domain architectures and genomic context. Consistent with the synteny observed in *Listeria* genomes, we found that the *inlP* domain architectures and genomic neighborhoods were highly conserved in *L. ivanovii londoniensis* and *L. seeligeri*, but not in *L. ivanovii ivanovii, L. innocua*, or *L. costaricensis*. While it is possible that these species could encode *inlP* homologs elsewhere in their genomes, it seems unlikely considering their domain architectures and lower conservation in sequence compared to other homologs. It is more likely that the homologous proteins identified in *Lc* and *Lin* are independent of InlP, but in the same class of small, secreted internalins that encompasses InlP, InlC, and InlH, among others^11^.

Notably, we found that *L. ivanovii londoniensis* encoded a homolog for *inlP* while *L. ivanovii ivanovii* did not. Historically, *L. ivanovii londoniensis* and *L. ivanovii ivanovii* have been distinguished biochemically^28^. Recently, Hupfeld *et al*., noted that the two subspecies could also be distinguished based on bacteriophage susceptibility: *L. ivanovii ivanovii* strains are sensitive to bacteriophages, while *L. ivanovii londoniensis* strains encode a type II-A CRISPR-Cas system rendering them resistant to many phages^29^. Our finding that only *L. ivanovii londoniensis*, and not *L. ivanovii ivanovii*, encodes the *inlP* gene provides another avenue for distinguishing between these two subspecies and could be beneficial to public health laboratories seeking to differentiate between them among clinical and food isolates. Additionally, because the evolution of virulence factors in *Listeria* remains mysterious, the specific presence of *inlP* in subspecies such as *ivanovii londoniensis* and its absence in *ivanovii ivanovii* may provide clues as to how *Listeria* evolves the ability to infect different cells and tissues.

Since *L. ivanovii* has been implicated in human and animal placental infection, it was not entirely surprising to find that it encoded the gene for InlP, an internalin known to enhance placental colonization. It was surprising, however, to find three copies of the *inlP* gene in *L. seeligeri* since it has not been significantly indicated in human or animal pathogenesis^30^. Our analyses suggested that InlP_*Ls2*_ was the most similar to InlP_*Lm*_ and InlP_*Liv*_. It is possible that this copy (InlP_*Ls2*_) is the ancestral one, and InlP_*Ls1*_ and InlP_*Ls3*_ resulted from subsequent duplication events. The presence of multiple homologs suggests that InlP could have alternative functions apart from enhancing placental colonization. *Listeria* species are frequently found in environmental isolates, as they readily reside in soil. It is possible that InlP provides a fitness advantage in this environment.

One of the main questions resulting from the discovery of InlP homologs centers on the evolutionary timeline of the *Listeria* genus: which InlP came first? Our discovery that InlP_*Liv*_, InlP_*Ls1*_, InlP_*Ls2*_, and InlP_*Ls3*_ do not contain the full-length LRR6 and LRR7 domains found in InlP_*Lm*_ begins to offer potential answers to this question. It is plausible that *L. monocytogenes, L. seeligeri*, and *L. ivanovii londoniensis* shared a common ancestor that passed down the *inlP* gene, and a subsequent insertion event in *L. monocytogenes* led to the full-length InlP containing LRR6 and LRR7. Conversely, it is likely that *L. monocytogenes* carries the ancestral copy of InlP (InlP_*Lm*_); the full-length *inlP* gene could have undergone a deletion resulting in the loss of LRR6 and LRR7 in InlP_*Liv*_ and InlP_*Ls*_, although it is less likely to observe several deletion events as against a single insertion event. Future studies on the evolution of the *Listeria* genus and the larger family of internalin proteins will be required to answer this question more rigorously and determine their possible links to pathogenicity.

An additional structural difference noted between newly identified InlP homologs resides in the Ca^2+^-binding loop of LRR3. Previously, this loop has been hypothesized to play a role in InlP signaling, activation, or stabilization in complex with its binding partner afadin^10^. Structural heterogeneity is visible in the Ca^2+^ regions of InlP homolog models; InlP homologs in *Liv* and *Ls* appear to have more open loops compared to InlP_*Lm*._ The ability of these loops to bind calcium, and their relative binding affinities will be an important avenue for future investigation, especially as more details regarding the function and regulation of InlP_*Lm*_ come to light.

Recent studies made several fundamental discoveries regarding the physical and mechanistic properties of InlP_*Lm*_ and its activity in the placenta, but many questions remain unanswered^9,10^. The discovery of InlP homologs in *L. ivanovii londoniensis* and *L. seeligeri*, two species that have not been substantially implicated in cases of placental infection, is compelling. Future studies will investigate the activity of these homologs to determine if they bind afadin and if they are able to enhance placental colonization of *L. monocytogenes* as well as endogenous InlP. Further, structural differences between these homologs suggest potential binding sites for the InlP-afadin interaction, which has not been resolved to date.

In summary, we report that *L. ivanovii londoniensis* and *L. seeligeri* encode homologs for the *L. monocytogenes* virulence factor InlP. Identified homologs in all three species are housed within similar genomic neighborhoods, flanked by the same housekeeping genes upstream and downstream; further, *L. seeligeri* encodes three copies of the *inlP* gene in this region. All four homologs are similar (>70%) to InlP in *L. monocytogenes*, the main structural difference resulting from the lack of full-length LRR6 and LRR7 regions in InlP_*Li*_, InlP_*Ls1*_, InlP_*Ls2*_, and InlP_*Ls3*_. Our findings will serve as a springboard for future evolutionary studies of internalins in the *Listeria* genus and will bolster future *in vitro* and *in vivo* studies of InlP in the context of virulence and pathogenicity.

## Author contributions

*Conceptualization:* KNC, JTB, JR, JWH. *Methodology:* KNC, JTB, JR. *Software:* JTB, JR. *Validation:* KNC, JTB, JR, JWH. *Formal Analysis:* KNC, JTB, JR. *Investigation:* KNC, JTB, JR. *Resources:* JR, JWH. *Data Curation:* KNC, JTB, JR. *Writing:* KNC, JR, JWH. *Visualization:* KNC, JTB, JR. *Supervision:* JR, JWH. *Project Administration:* JR, JWH. *Funding:* JR, JWH.

## Data availability and reuse

All the data, analyses, and visualizations are available in our GitHub repository, https://github.com/JRaviLab/inlp_listeria. Text, figures, and data are licensed under Creative Commons Attribution CC BY 4.0.

## Supplementary Figures

**Figure S1:**
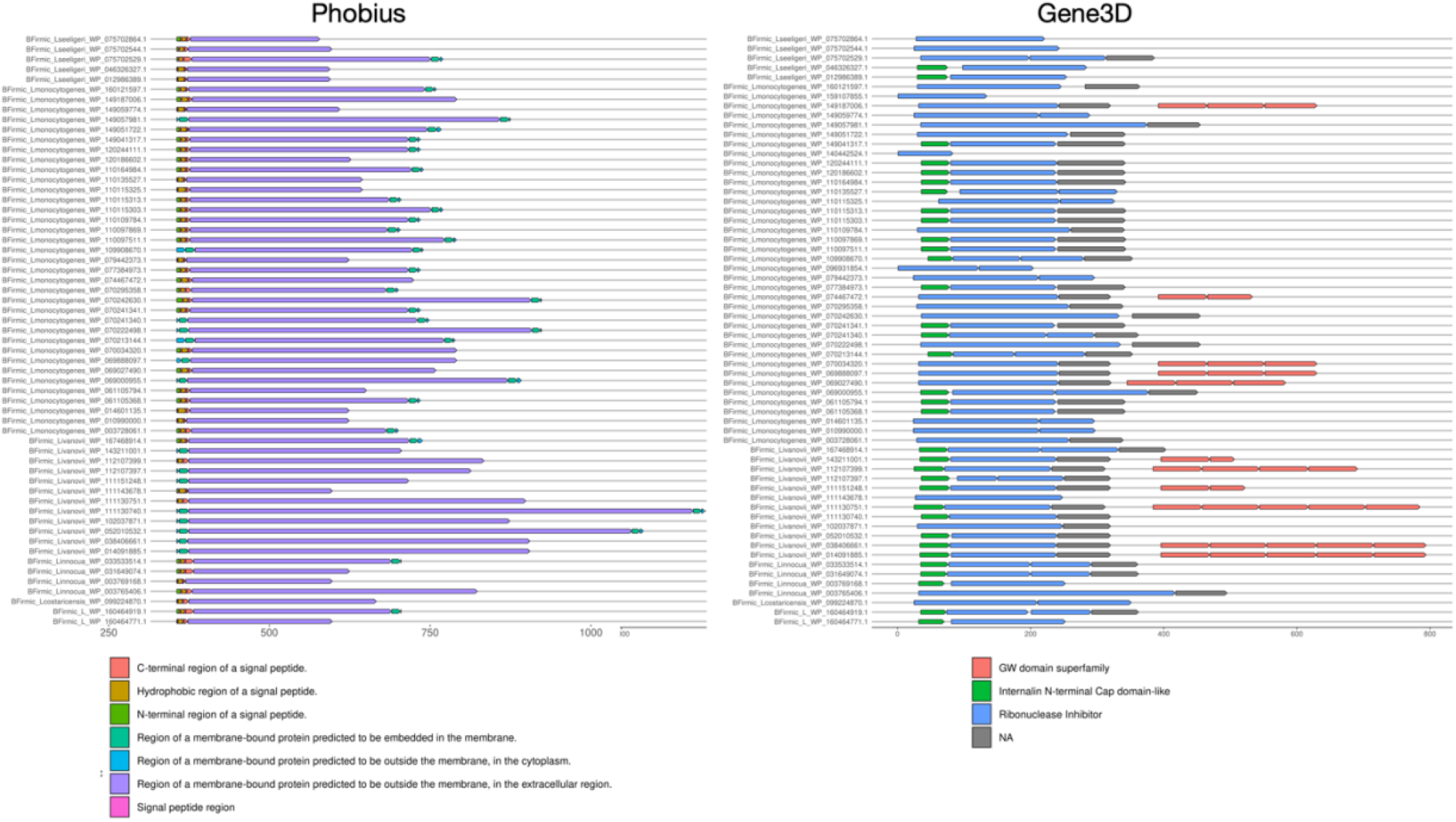
Phobius and Gene3D Domain Architectures of Identified Internalin P Homologs. Cellular localizations (Phobius) and domain architectures (Gene3D) were determined and visualized using MolEvolvR. Figure legends correspond to different domain predictions.

**Figure S2:**
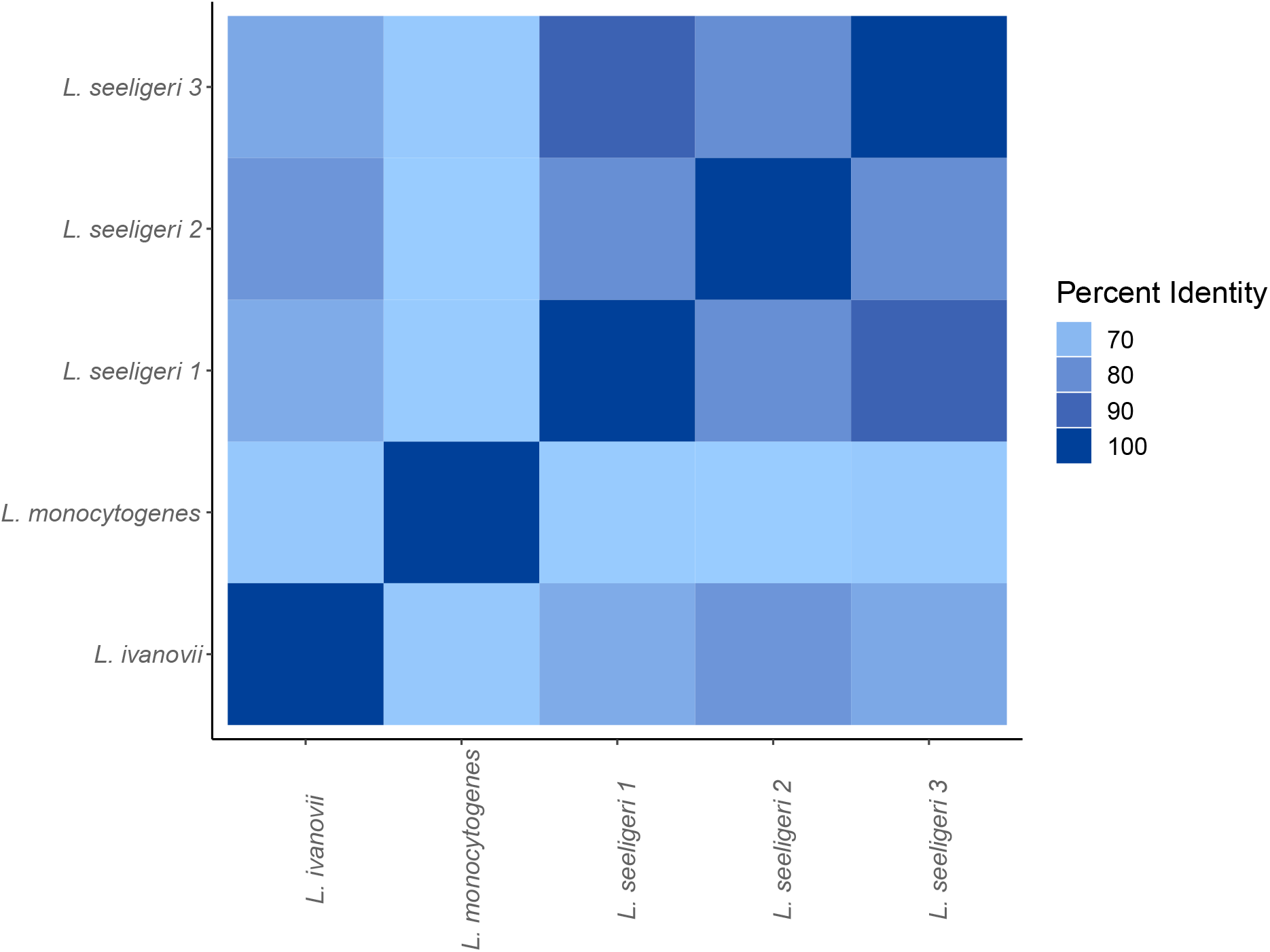
Percent Nucleotide Identity of InlP Homolog Genes in *L. monocytogenes, L. ivanovii londoniensis*, and *L. seeligeri*. Heatmap representing pairwise percent identities of the nucleotide sequences of Internalin P homologs in *L. monocytogenes, L. ivanovii londoniensis*, and *L. seeligeri*. Darker and lighter shades of blue represent higher and lower percent identities, respectively.

