## Supplemental Table 1 for "Novel Internalin P homologs in *Listeria*"

| Query | Subject | Species | Lineage | Accession | PcPositive | PcIdentity | DomArch.Pfam |
|---|---|---|---|---|---|---|---|
| BFirmic_Livanovii_WP_111143678.1 | BFirmic_Livanovii_WP_111143678.1 | Listeria ivanovii | Bacteria>Firmicutes | WP_111143678.1 | 100 | 99.71 | Internalin_N |
| BFirmic_Livanovii_WP_111143678.1 | BFirmic_Linnocua_WP_003769168.1 | Listeria innocua | Bacteria>Firmicutes | WP_003769168.1 | 60.29 | 37.02 | Internalin_N |
| BFirmic_Livanovii_WP_111143678.1 | BFirmic_L_WP_160464771.1 | Listeria | Bacteria>Firmicutes | WP_160464771.1 | 60.29 | 36.46 | Internalin_N |
| BFirmic_Livanovii_WP_111143678.1 | BFirmic_Lseeligeri_WP_075702864.1 | Listeria seeligeri | Bacteria>Firmicutes | WP_075702864.1 | 41.43 | 28.62 | NA |
| BFirmic_Livanovii_WP_111143678.1 | BFirmic_Lmonocytogenes_WP_077384973.1 | Listeria monocytogenes | Bacteria>Firmicutes | WP_077384973.1 | 40.86 | 22.29 | Internalin_N+<br>LRR_8+<br>LRR_adjacent+<br>Flg_new+<br>Flg_new |
| BFirmic_Livanovii_WP_111143678.1 | BFirmic_Lmonocytogenes_WP_061105368.1 | Listeria monocytogenes | Bacteria>Firmicutes | WP_061105368.1 | 40.58 | 23.36 | Internalin_N+<br>LRR_4+<br>LRR_adjacent+<br>Flg_new+<br>Flg_new |
| BFirmic_Livanovii_WP_111143678.1 | BFirmic_Lmonocytogenes_WP_070241341.1 | Listeria monocytogenes | Bacteria>Firmicutes | WP_070241341.1 | 40.57 | 22.57 | Internalin_N+<br>LRR_4+<br>LRR_6+<br>LRR_4+<br>LRR_adjacent+<br>Flg_new+<br>Flg_new |
| BFirmic_Livanovii_WP_111143678.1 | BFirmic_Lmonocytogenes_WP_120186602.1 | Listeria monocytogenes | Bacteria>Firmicutes | WP_120186602.1 | 40.57 | 22.29 | Internalin_N+<br>LRR_6+<br>LRR_4+<br>LRR_4+<br>LRR_adjacent+<br>Flg_new |
| BFirmic_Livanovii_WP_111143678.1 | BFirmic_Lmonocytogenes_WP_061105794.1 | Listeria monocytogenes | Bacteria>Firmicutes | WP_061105794.1 | 40.29 | 22 | Internalin_N+<br>LRR_4+<br>LRR_4+<br>LRR_adjacent+<br>Flg_new |
| BFirmic_Livanovii_WP_111143678.1 | BFirmic_Lmonocytogenes_WP_110097511.1 | Listeria monocytogenes | Bacteria>Firmicutes | WP_110097511.1 | 40.29 | 22.29 | Internalin_N+<br>LRR_4+<br>LRR_4+<br>LRR_adjacent+<br>LRR_adjacent+<br>Flg_new+<br>Flg_new+<br>Flg_new |
| BFirmic_Livanovii_WP_111143678.1 | BFirmic_Lmonocytogenes_WP_110097869.1 | Listeria monocytogenes | Bacteria>Firmicutes | WP_110097869.1 | 40.29 | 22.29 | Internalin_N+<br>LRR_4+<br>LRR_4+<br>LRR_adjacent+<br>LRR_adjacent+<br>Flg_new+<br>Flg_new |
| BFirmic_Livanovii_WP_111143678.1 | BFirmic_Lmonocytogenes_WP_110115303.1 | Listeria monocytogenes | Bacteria>Firmicutes | WP_110115303.1 | 40.29 | 22.29 | Internalin_N+<br>LRR_4+<br>LRR_4+<br>LRR_adjacent+<br>Flg_new+<br>Flg_new+<br>Flg_new |
| BFirmic_Livanovii_WP_111143678.1 | BFirmic_Lmonocytogenes_WP_110115313.1 | Listeria monocytogenes | Bacteria>Firmicutes | WP_110115313.1 | 40.29 | 22.29 | Internalin_N+<br>LRR_4+<br>LRR_4+<br>LRR_adjacent+<br>LRR_adjacent+<br>Flg_new |
| BFirmic_Livanovii_WP_111143678.1 | BFirmic_Lmonocytogenes_WP_149041317.1 | Listeria monocytogenes | Bacteria>Firmicutes | WP_149041317.1 | 40.29 | 22.57 | Internalin_N+<br>LRR_4+<br>LRR_6+<br>LRR_4+<br>LRR_4+<br>LRR_adjacent+<br>Flg_new+<br>Flg_new |
| BFirmic_Livanovii_WP_111143678.1 | BFirmic_Lmonocytogenes_WP_070034320.1 | Listeria monocytogenes | Bacteria>Firmicutes | WP_070034320.1 | 40.28 | 21.91 | Internalin_N+<br>LRR_4+<br>LRR_adjacent+<br>Flg_new+<br>GW+<br>GW+<br>GW |
| BFirmic_Livanovii_WP_111143678.1 | BFirmic_Lmonocytogenes_WP_149187006.1 | Listeria monocytogenes | Bacteria>Firmicutes | WP_149187006.1 | 40.28 | 22.22 | Internalin_N+<br>LRR_8+<br>LRR_4+<br>LRR_adjacent+<br>Flg_new+<br>GW+<br>GW+<br>GW |
| BFirmic_Livanovii_WP_111143678.1 | BFirmic_Livanovii_WP_052010532.1 | Listeria ivanovii | Bacteria>Firmicutes | WP_052010532.1 | 39.72 | 21.75 | Internalin_N+<br>LRR_4+<br>LRR_adjacent+<br>Flg_new+<br>Flg_new+<br>Flg_new+<br>Flg_new+<br>Flg_new+<br>Flg_new+<br>Flg_new+<br>Flg_new+<br>Flg_new+<br>Gram_pos_anchor |
| BFirmic_Livanovii_WP_111143678.1 | BFirmic_Lmonocytogenes_WP_110164984.1 | Listeria monocytogenes | Bacteria>Firmicutes | WP_110164984.1 | 39.71 | 22.57 | Internalin_N+<br>LRR_6+<br>LRR_4+<br>LRR_4+<br>LRR_adjacent+<br>LRR_adjacent+<br>Flg_new+<br>Flg_new |
| BFirmic_Livanovii_WP_111143678.1 | BFirmic_Livanovii_WP_102037871.1 | Listeria ivanovii | Bacteria>Firmicutes | WP_102037871.1 | 38.86 | 22.06 | Internalin_N+<br>LRR_4+<br>LRR_adjacent+<br>Flg_new+<br>Flg_new+<br>Flg_new+<br>Flg_new+<br>Flg_new+<br>Flg_new |
| BFirmic_Livanovii_WP_111143678.1 | BFirmic_Livanovii_WP_111130740.1 | Listeria ivanovii | Bacteria>Firmicutes | WP_111130740.1 | 38.86 | 22.06 | Internalin_N+<br>LRR_4+<br>LRR_adjacent+<br>Flg_new+<br>Flg_new+<br>Flg_new+<br>Flg_new+<br>Flg_new+<br>Flg_new+<br>Flg_new+<br>Flg_new+<br>Flg_new+<br>Flg_new+<br>Flg_new+<br>Gram_pos_anchor |
| BFirmic_Livanovii_WP_111143678.1 | BFirmic_Livanovii_WP_112107397.1 | Listeria ivanovii | Bacteria>Firmicutes | WP_112107397.1 | 38.86 | 22.06 | Internalin_N+<br>LRR_4+<br>LRR_adjacent+<br>Flg_new+<br>Flg_new+<br>Flg_new+<br>Flg_new+<br>Flg_new |
| BFirmic_Livanovii_WP_111143678.1 | BFirmic_Lmonocytogenes_WP_069888097.1 | Listeria monocytogenes | Bacteria>Firmicutes | WP_069888097.1 | 37.71 | 21.27 | Internalin_N+<br>LRR_8+<br>LRR_adjacent+<br>Flg_new+<br>GW+<br>GW+<br>GW |
| BFirmic_Livanovii_WP_111143678.1 | BFirmic_Livanovii_WP_014091885.1 | Listeria ivanovii | Bacteria>Firmicutes | WP_014091885.1 | 37.43 | 23.68 | Internalin_N+<br>LRR_4+<br>LRR_adjacent+<br>Flg_new+<br>GW+<br>GW+<br>GW+<br>GW+<br>GW |
| BFirmic_Livanovii_WP_111143678.1 | BFirmic_Livanovii_WP_111151248.1 | Listeria ivanovii | Bacteria>Firmicutes | WP_111151248.1 | 37.43 | 23.68 | Internalin_N+<br>LRR_4+<br>LRR_adjacent+<br>Flg_new+<br>GW+<br>GW |
| BFirmic_Livanovii_WP_111143678.1 | BFirmic_Livanovii_WP_143211001.1 | Listeria ivanovii | Bacteria>Firmicutes | WP_143211001.1 | 37.43 | 23.68 | Internalin_N+<br>LRR_4+<br>LRR_adjacent+<br>Flg_new |
| BFirmic_Livanovii_WP_111143678.1 | BFirmic_Lmonocytogenes_WP_069027490.1 | Listeria monocytogenes | Bacteria>Firmicutes | WP_069027490.1 | 36.85 | 20.95 | Internalin_N+<br>LRR_8+<br>LRR_4+<br>LRR_adjacent+<br>GW+<br>GW+<br>GW |
| BFirmic_Livanovii_WP_111143678.1 | BFirmic_Lmonocytogenes_WP_074467472.1 | Listeria monocytogenes | Bacteria>Firmicutes | WP_074467472.1 | 36.85 | 20.95 | Internalin_N+<br>LRR_8+<br>LRR_4+<br>LRR_adjacent+<br>Flg_new+<br>GW+<br>GW |
| BFirmic_Livanovii_WP_111143678.1 | BFirmic_Lmonocytogenes_WP_070295358.1 | Listeria monocytogenes | Bacteria>Firmicutes | WP_070295358.1 | 36 | 21.93 | Internalin_N+<br>LRR_4+<br>LRR_adjacent+<br>Flg_new |
| BFirmic_Livanovii_WP_111143678.1 | BFirmic_Livanovii_WP_111130751.1 | Listeria ivanovii | Bacteria>Firmicutes | WP_111130751.1 | 34.29 | 20.2 | Internalin_N+<br>LRR_8+<br>LRR_adjacent+<br>Flg_new+<br>GW+<br>GW+<br>GW+<br>GW+<br>GW |
| BFirmic_Livanovii_WP_111143678.1 | BFirmic_Livanovii_WP_112107399.1 | Listeria ivanovii | Bacteria>Firmicutes | WP_112107399.1 | 34.29 | 20.2 | Internalin_N+<br>LRR_8+<br>LRR_adjacent+<br>Flg_new+<br>GW+<br>GW+<br>GW+<br>GW |
| BFirmic_Livanovii_WP_111143678.1 | BFirmic_Lmonocytogenes_WP_109908670.1 | Listeria monocytogenes | Bacteria>Firmicutes | WP_109908670.1 | 31.43 | 24.81 | Internalin_N+<br>LRR_4+<br>LRR_8+<br>LRR_adjacent+<br>Flg_new+<br>Flg_new |
| monocytogenes_inlP | BFirmic_Lmonocytogenes_WP_014601135.1 | Listeria monocytogenes | Bacteria>Firmicutes | WP_014601135.1 | 100 | 99.74 | NA |
| monocytogenes_inlP | BFirmic_Lmonocytogenes_WP_010990000.1 | Listeria monocytogenes | Bacteria>Firmicutes | WP_010990000.1 | 99.74 | 99.48 | Internalin_N+<br>LRR_4+<br>LRR_1 |
| monocytogenes_inlP | BFirmic_Lmonocytogenes_WP_110115325.1 | Listeria monocytogenes | Bacteria>Firmicutes | WP_110115325.1 | 99.49 | 92.11 | Internalin_N+<br>Internalin_N+<br>LRR_4+<br>LRR_1 |
| monocytogenes_inlP | BFirmic_Lmonocytogenes_WP_110135527.1 | Listeria monocytogenes | Bacteria>Firmicutes | WP_110135527.1 | 99.49 | 91.63 | Internalin_N+<br>LRR_8 |
| monocytogenes_inlP | BFirmic_Lmonocytogenes_WP_079442373.1 | Listeria monocytogenes | Bacteria>Firmicutes | WP_079442373.1 | 99.23 | 98.45 | Internalin_N+<br>LRR_4 |
| monocytogenes_inlP | BFirmic_Lmonocytogenes_WP_096931854.1 | Listeria monocytogenes | Bacteria>Firmicutes | WP_096931854.1 | 74.42 | 99.65 | LRR_4+<br>LRR_1 |
| monocytogenes_inlP | BFirmic_Lmonocytogenes_WP_159107855.1 | Listeria monocytogenes | Bacteria>Firmicutes | WP_159107855.1 | 58.14 | 93.1 | LRR_8+<br>LRR_8 |
| monocytogenes_inlP | BFirmic_Lcostaricensis_WP_099224870.1 | Listeria costaricensis | Bacteria>Firmicutes | WP_099224870.1 | 55.04 | 31 | NA |
| monocytogenes_inlP | BFirmic_Lmonocytogenes_WP_140442524.1 | Listeria monocytogenes | Bacteria>Firmicutes | WP_140442524.1 | 36.95 | 91.28 | LRR_1 |
| monocytogenes_inlP | BFirmic_Lmonocytogenes_WP_070242630.1 | Listeria monocytogenes | Bacteria>Firmicutes | WP_070242630.1 | 34.63 | 20.41 | Internalin_N+<br>LRR_6+<br>LRR_4+<br>LRR_8+<br>LRR_6+<br>LRR_adjacent+<br>Flg_new+<br>Flg_new+<br>Flg_new+<br>Flg_new |
| monocytogenes_inlP | BFirmic_Lmonocytogenes_WP_070241340.1 | Listeria monocytogenes | Bacteria>Firmicutes | WP_070241340.1 | 32.56 | 16.08 | Internalin_N+<br>LRR_adjacent+<br>Flg_new+<br>Flg_new |
| monocytogenes_inlP | BFirmic_Linnocua_WP_003770165.1 | Listeria innocua | Bacteria>Firmicutes | WP_003770165.1 | 31.78 | 20.51 | NA |
| monocytogenes_inlP | BFirmic_Linnocua_WP_033533514.1 | Listeria innocua | Bacteria>Firmicutes | WP_033533514.1 | 31.78 | 20.19 | Internalin_N+<br>LRR_4+<br>LRR_adjacent+<br>Flg_new |
| monocytogenes_inlP | BFirmic_L_WP_045554046.1 | Listeria | Bacteria>Firmicutes | WP_045554046.1 | 31.78 | 20.19 | NA |
| monocytogenes_inlP | BFirmic_L_WP_160464919.1 | Listeria | Bacteria>Firmicutes | WP_160464919.1 | 31.52 | 19.87 | Internalin_N+<br>LRR_4+<br>LRR_adjacent+<br>Flg_new |
| monocytogenes_inlP | BFirmic_Linnocua_WP_031649074.1 | Listeria innocua | Bacteria>Firmicutes | WP_031649074.1 | 31.27 | 20.2 | Internalin_N+<br>LRR_4+<br>LRR_adjacent |
| monocytogenes_inlP | BFirmic_Lmonocytogenes_WP_070222498.1 | Listeria monocytogenes | Bacteria>Firmicutes | WP_070222498.1 | 31.27 | 22.36 | Internalin_N+<br>LRR_4+<br>LRR_6+<br>LRR_8+<br>LRR_6+<br>LRR_adjacent+<br>Flg_new+<br>Flg_new+<br>Flg_new+<br>Flg_new |
| monocytogenes_inlP | BFirmic_Lmonocytogenes_WP_149057981.1 | Listeria monocytogenes | Bacteria>Firmicutes | WP_149057981.1 | 31.27 | 22.36 | Internalin_N+<br>LRR_4+<br>LRR_6+<br>LRR_8+<br>LRR_6+<br>LRR_adjacent+<br>Flg_new+<br>Flg_new+<br>Flg_new |
| monocytogenes_inlP | BFirmic_Livanovii_WP_167468914.1 | Listeria ivanovii | Bacteria>Firmicutes | WP_167468914.1 | 30.23 | 17.06 | Internalin_N+<br>LRR_4+<br>LRR_6+<br>LRR_4+<br>LRR_adjacent+<br>Flg_new |
| monocytogenes_inlP | BFirmic_Lseeligeri_WP_075702529.1 | Listeria seeligeri | Bacteria>Firmicutes | WP_075702529.1 | 29.72 | 18.91 | Internalin_N+<br>LRR_4+<br>LRR_4+<br>LRR_adjacent+<br>Flg_new+<br>Flg_new+<br>Gram_pos_anchor |
| monocytogenes_inlP | BFirmic_Linnocua_WP_003765406.1 | Listeria innocua | Bacteria>Firmicutes | WP_003765406.1 | 29.71 | 19.15 | Internalin_N+<br>LRR_4+<br>LRR_4+<br>LRR_4+<br>LRR_4+<br>LRR_adjacent+<br>Flg_new+<br>Flg_new+<br>Flg_new |
| BFirmic_Lseeligeri_WP_012986389.1 | BFirmic_Lseeligeri_WP_012986389.1 | Listeria seeligeri | Bacteria>Firmicutes | WP_012986389.1 | 100 | 100 | Internalin_N |
| BFirmic_Lseeligeri_WP_012986389.1 | BFirmic_Lseeligeri_WP_046326327.1 | Listeria seeligeri | Bacteria>Firmicutes | WP_046326327.1 | 94.51 | 87.57 | Internalin_N+<br>LRR_8 |
| BFirmic_Lseeligeri_WP_012986389.1 | BFirmic_Lmonocytogenes_WP_149051722.1 | Listeria monocytogenes | Bacteria>Firmicutes | WP_149051722.1 | 41.32 | 21.02 | Internalin_N+<br>LRR_4+<br>LRR_4+<br>LRR_4+<br>LRR_adjacent+<br>Flg_new+<br>Flg_new |
| seeligeri_2_RS12045 | BFirmic_Lmonocytogenes_WP_149059774.1 | Listeria monocytogenes | Bacteria>Firmicutes | WP_149059774.1 | 81.85 | 64.6 | Internalin_N |
| seeligeri_2_RS12045 | BFirmic_Livanovii_WP_038406661.1 | Listeria ivanovii | Bacteria>Firmicutes | WP_038406661.1 | 36.6 | 20.74 | NA |
| seeligeri_2_RS12045 | BFirmic_Lmonocytogenes_WP_160121597.1 | Listeria monocytogenes | Bacteria>Firmicutes | WP_160121597.1 | 29.97 | 18.22 | Internalin_N+<br>LRR_4+<br>LRR_4+<br>LRR_adjacent+<br>Flg_new+<br>Flg_new+<br>Gram_pos_anchor |
| seeligeri_3_RS12050 | BFirmic_Lseeligeri_WP_075702544.1 | Listeria seeligeri | Bacteria>Firmicutes | WP_075702544.1 | 59.82 | 38.94 | LRR_8 |
| seeligeri_3_RS12050 | BFirmic_Lmonocytogenes_WP_120244111.1 | Listeria monocytogenes | Bacteria>Firmicutes | WP_120244111.1 | 41.91 | 21.35 | Internalin_N+<br>LRR_4+<br>LRR_4+<br>LRR_adjacent+<br>Flg_new+<br>Flg_new |
| seeligeri_3_RS12050 | BFirmic_Lmonocytogenes_WP_110109784.1 | Listeria monocytogenes | Bacteria>Firmicutes | WP_110109784.1 | 40.75 | 20.84 | Internalin_N+<br>LRR_6+<br>LRR_4+<br>LRR_4+<br>LRR_adjacent+<br>Flg_new+<br>Flg_new |
| seeligeri_3_RS12050 | BFirmic_Lmonocytogenes_WP_003728061.1 | Listeria monocytogenes | Bacteria>Firmicutes | WP_003728061.1 | 39.88 | 21.07 | Internalin_N+<br>LRR_4+<br>LRR_4+<br>LRR_4+<br>LRR_adjacent+<br>Flg_new |
| seeligeri_3_RS12050 | BFirmic_Lmonocytogenes_WP_069000955.1 | Listeria monocytogenes | Bacteria>Firmicutes | WP_069000955.1 | 32.95 | 18.05 | Internalin_N+<br>LRR_4+<br>LRR_4+<br>LRR_4+<br>LRR_adjacent+<br>Flg_new+<br>Flg_new+<br>Flg_new |
| seeligeri_3_RS12050 | BFirmic_Lmonocytogenes_WP_070213144.1 | Listeria monocytogenes | Bacteria>Firmicutes | WP_070213144.1 | 28.9 | 22.35 | Internalin_N+<br>LRR_4+<br>LRR_8+<br>LRR_adjacent+<br>Flg_new+<br>Flg_new+<br>Flg_new |

More information available on GitHub.
